## Supplementary Information for "A new model for the HPA axis explains dysregulation of stress hormones on the timescale of weeks"

Karin et al.

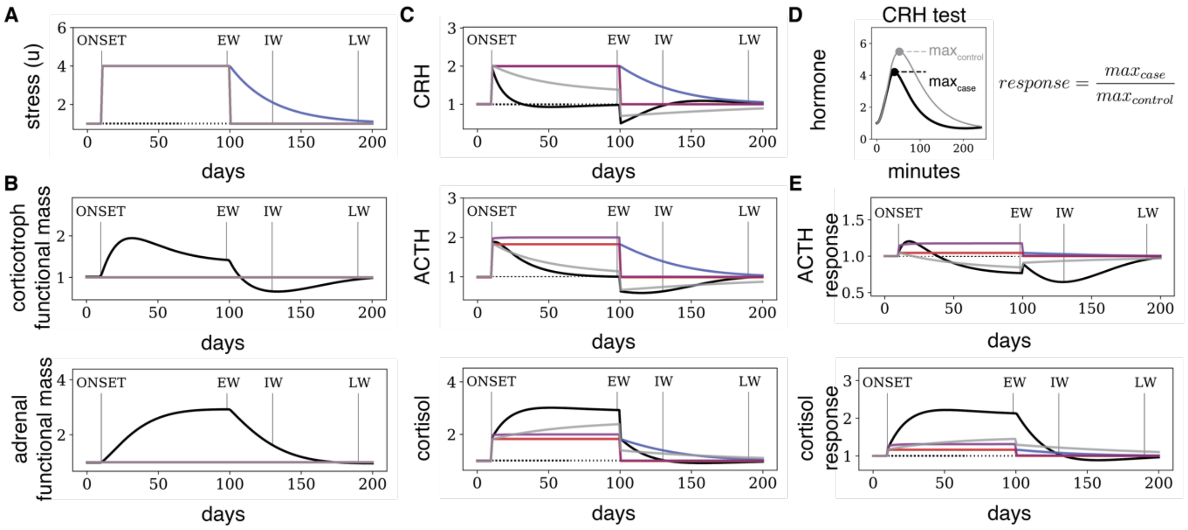

Figure S1. Model dynamics of HPA axis after prolonged stress. Numerical solutions of the HPA models after a prolonged pulse of stress input, as in Figure 3 of the main text. The model with gland functional mass dynamics (black lines) recapitulates the dynamics observed in Figure 1. Models in which mass is constant (red lines) or models with constant masses and (i) month-timescale drop in input signal (blue lines), (ii) acquisition of glucocorticoid resistance (purple lines), do not develop a blunting of ACTH responses or a mismatch between ACTH and cortisol dynamics. A model with (iii) desensitization of cortisol clearance rate (gray line) does develop a blunting of ACTH responses, but does not recapitulate the mismatch observation that ACTH dynamics normalize after cortisol dynamics.

### Supplementary Information 1. Several alternative models for slow timescale dynamics do not explain observed mismatch between ACTH and cortisol.

In this section we tested whether alternative models can explain the observations at the focus of this study, presented in Figure 1: blunting of ACTH responses to CRH challenges after prolonged activation of the HPA axis, which persists for weeks-months after cortisol levels and dynamics normalize. Since this persistence occurs on the timescale of weeks-months, it cannot be explained by standard hormone dynamics with a timescale of hours (red lines in Figure S1). We thus explored alternative models without gland mass dynamics, but with a relevant timescale on the order of weeks-months. The models are described in the Methods section. The model dynamics are depicted with colored lines in Figure S1, with the mass-change model of the main text in thick black line for comparison. None of the alternative models has mass changes so that the dynamics in Figure S1B,C are flat, and all lines overlap.

The first alternative model that we explored is a model where the input signal drops gradually (Methods, blue line in Figure S1A). In this model, prolonged activation of the HPA axis does not result in blunted ACTH responses, but instead in elevated ACTH levels and responses (Figure S1CE, blue lines). Moreover, the recovery of ACTH and cortisol is coordinated in this model and there is no mismatch.

A second alternative model is glucocorticoid resistance which depends on cortisol levels, a mechanism which has some empirical support (Cohen *et al*, 2012; Merkulov *et al*, 2017). We model this mechanism by adding a cortisol-dependent decrease in the sensitivity of the GR (Methods, purple lines in Figure

S1). This mechanism also does not result in blunted ACTH responses (in fact, it sensitizes ACTH responses), and there is no mismatch between cortisol and ACTH after recovery from prolonged stress. Finally, we considered a mechanism that can reproduce persistent blunting of ACTH – a putative cortisol-dependent drop in cortisol clearance rate (Methods, gray lines in Figure S1). While this mechanism reproduces both persistent blunting and hypercortisolemia after chronic stress (Figure S1E gray lines), it does not show a mismatch between the recovery of cortisol and that of ACTH (they both recover together), and therefore it cannot explain the data of Figure 1.

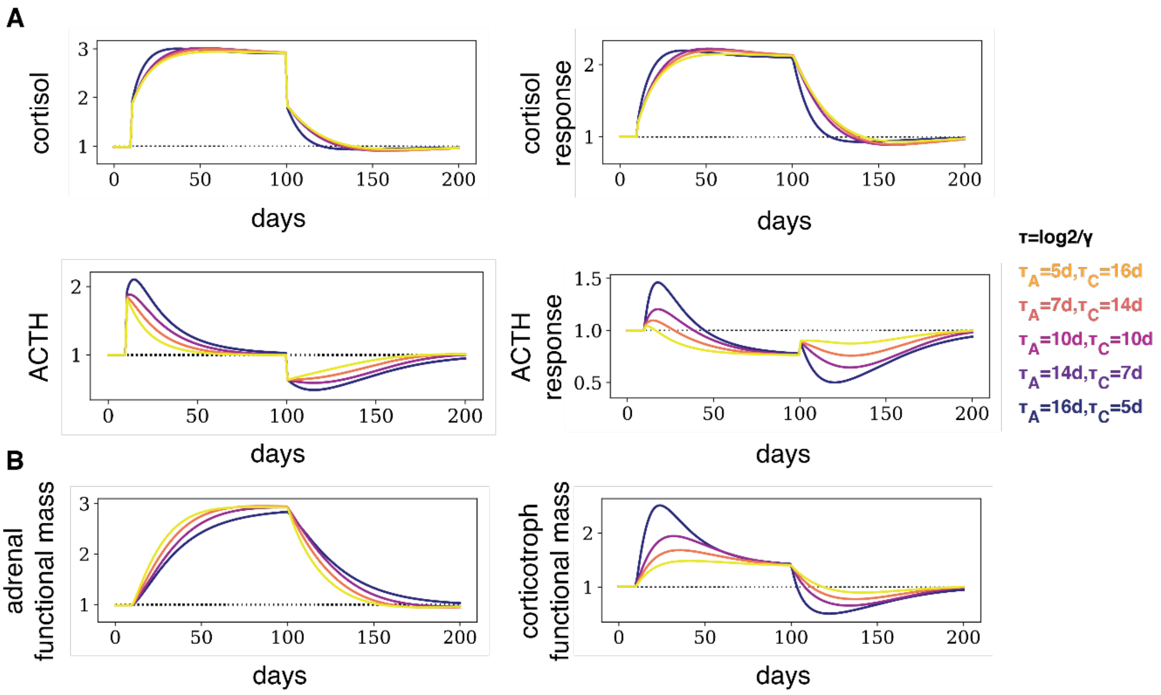

Figure S2. Model dynamics for various turnover rates of pituitary corticotrophs and adrenal cortex cells. (AB) Here we show the dynamics of the HPA axis during and after a prolonged pulse of stress, described in Figure 3, for various values of $\tau_A$ (adrenal half-life) and $\gamma_C$ (corticotroph turnover). The faster the turnover of the corticotrophs relative to the turnover of the adrenal cortex, the more pronounced is the dysregulation of ACTH dynamics after prolonged stress. On the other hand, a faster turnover of corticotrophs also causes a faster return to baseline of cortisol levels and responses. For all parameter values, however, ACTH always recovers after cortisol and CRH after prolonged stress.

### Supplementary Information 2. Dependence of recovery from prolonged stress on corticotroph and adrenal turnover rate parameters.

We tested whether the qualitative features we discuss are robust to changes in model parameters. We note that all fast-timescale parameters (namely the hormone removal rates $w_1$,$w_2$,$w_3$ and specific production rates $k_1$,$k_2$,$k_3$) do not affect the slow timescale of weeks. The two parameters that do affect the weeks-timescale are the *turnover parameters* of the functional masses of the corticotrophs and the adrenal cortex, $w_C, w$. To illustrate the dependence on these two parameters, we plotted in Figure S2 the dynamics of the HPA axis variables with different values of the parameters. For ease of interpretation, we present this in terms of tissue half-lives ($\tau_C = \frac{\log 2}{w_C}, \tau_A = \frac{\log 2}{w_A}$). We find that for half-lives in the range of a few days to a few weeks, the dynamics of the HPA axis show the same qualitative features that we discussed: a return to baseline cortisol levels and CRH-test responses within a few weeks, and a return to baseline of ACTH level and responses only later, within a few months. This holds regardless of whether the corticotrophs have a faster or slower half-life than the adrenal cortex. However, the faster the turnover of the corticotrophs is compared with that of the adrenal cortex ($\tau_C \ll \tau_A$), the more severe is the blunting of ACTH in the weeks following the cessation of the stressor. On the other hand, a faster turnover of corticotrophs also leads to faster recovery of cortisol, suggesting a tradeoff between faster recovery of cortisol levels with milder blunting of ACTH (and with it presumably other POMC peptides including endorphins with physiological consequences).

Supplementary Information 3. Analysis of the dynamical compensation properties of the HPA model.

Here we analyze the dynamical compensation properties of the general (un-normalized) equations of the HPA axis (Eq. 6-10 in Methods). The model equations are, writing $x_1$, $x_2$ and $x_3$ for CRH, ACTH and cortisol:

$$\frac{dx_1}{dt} = k_1 u(t) \cdot M(x_3) \cdot G(x_3) - w_1 x_1 \quad [1]$$

$$\frac{dx_2}{dt} = k_2 C x_1 \cdot G(x_3) - w_2 x_2 \quad [2]$$

$$\frac{dx_3}{dt} = k_3\, A x_3 - w_3 x_3 \quad [3]$$

$$\frac{dC}{dt} = C(k_C x_1 - w_c\,) \quad [4]$$

$$\frac{dA}{dt} = A(k_A x_2 - w_A) \quad [5]$$

The $w_i$ parameters are the removal rates, and the $k_i$ parameters are the hormone production rates. We note here that we focus on the slow timescale of weeks, and the rates $k_1, k_2, k_3$ do not affect the conclusions of the paper. By equating all derivatives to zero, the steady-state solution of the model is (for $x_3 \ll K_{GR}$):

$$x_{1,st} = \frac{w_P}{k_P}$$

$$x_{2,st} = \frac{w_A}{k_A}$$

$$x_{3,st} = \frac{k_1 k_P}{w_1 w_P} u$$

$$C_{st} = \frac{k_C w_2 \mathrm{w_A}}{w_C k_2 k_A}$$

$$A_{st} = \frac{w_3 k_A \frac{u k_1 k_C}{w_1 w_C}}{k_3 w_A}$$

We next define dynamical compensation and test it in this model. A system has dynamical compensation with respect to parameter $p$ and variable $x$, if for any input $u(t)$ and any constant value of $p$ the dynamics of the variable $x$ does not depends on $p$, provided the initial conditions are at steady-state (Karin *et al*, 2016). To rule out trivial cases, such as systems in which $p$ does not affect the dynamics at all or is non-identifiable (Karin *et al*, 2017), we further require that when the system is out of steady-state, the output $x(t)$ for a given $u(t)$ *does* depend on the value of $p$.

A necessary condition for dynamical compensation is that the *steady-state* value of the variable $x$ is independent of $p$. Thus, it is easy to rule out dynamical compensation when the steady state value depends on the parameter. To establish dynamical compensation for the rest of the cases, we used the method of Karin et al. (Karin *et al*, 2016) ,which employs transformed variables to prove invariance.

We find that in the HPA model many variables are dynamically compensated for the secretion rates $k_i$. For example, cortisol $x_3$ has dynamical compensation with respect to the secretion rates of $C$ and $A$, and the proliferation rate of $A$. No variable, however, has dynamical compensation with respect to the removal rates $w_i$ since they change the timescale of the dynamics.

| | $CRH$ $x_1$ | $ACTH$ $x_2$ | $CORT$ $x_3$ | $C$ | $A$ |
|---|---|---|---|---|---|
| $k_1$ | + | + | | | |
| $k_2$ | + | + | + | | + |
| $k_3$ | + | + | + | + | |
| $k_C$ | | + | | | |
| $k_A$ | + | | + | | |
| $w_C$ | | + | | | |
| $w_A$ | + | | + | | |

**Table S1.** Dynamical compensation in the HPA model for the case of $CORT \ll K_{GR}$. Compensated variables are in the columns and respective compensated parameters are in the rows. + marks represent pairs of dynamically compensated parameter and variable.

Note that the above table holds for the case where the GR is not activated ($CORT \ll K_{GR}$). In the more general case, where $CORT$ can be similar or larger than $K_{GR}$, the following table holds:

| | $CRH$ $x_1$ | $ACTH$ $x_2$ | $CORT$ $x_3$ | $C$ | $A$ |
|---|---|---|---|---|---|
| $k_2$ | + | + | + | | + |
| $k_3$ | + | + | + | + | |
| $k_A$ | + | | + | | |
| $w_A$ | + | | + | | |

**Table S2.** Dynamical compensation in the HPA model for the general case. Compensated variables are in the columns and respective compensated parameters are in the rows. + marks represent pairs of dynamically compensated parameter and variable.

Of special importance are the parameters $k_2$ and $k_3$, because they are expected to vary physiologically. These hormone secretion parameters (per unit biomass and per unit upstream hormone) can vary with the metabolic state of the corticotrophs and adrenal cells, the number of receptors for the upstream hormone, neuronal inputs, cytokines and other factors. They also vary with blood volume because increased blood volume dilutes out blood hormone levels and effectively changes these parameters (which is inversely proportional to blood volume) (Karin *et al*, 2016).

Thus, dynamical compensation ensures a proper cortisol and ACTH response regardless of physiological changes that affect the specific secretion rates $k_2$ and $k_3$. It ensures that a given stress input u(t) provides an output time course of ACTH and cortisol which does not depend on the secretion parameters, provided the initial conditions including C and A masses are at steady state. The phenomena described in the main text are transients in which C and A have not yet reached steady state, resulting in blunted hormone responses to CRH tests. Long after the prolonged stress described in Figure 2,3, the system returns to steady state and responses return to normal.

**References**.

Cohen S, Janicki-Deverts D, Doyle WJ, Miller GE, Frank E, Rabin BS & Turner RB (2012) Chronic stress, glucocorticoid receptor resistance, inflammation, and disease risk. *Proc. Natl. Acad. Sci.* **109:** 5995–5999

Karin O, Alon U & Sontag E (2017) A Note On Dynamical Compensation And Its Relation To Parameter Identifiability. *bioRxiv***:** 123489

Karin O, Swisa A, Glaser B, Dor Y & Alon U (2016) Dynamical compensation in physiological circuits. *Mol. Syst. Biol.* **12:** 886

Merkulov VM, Merkulova TI & Bondar NP (2017) Mechanisms of brain glucocorticoid resistance in stress-induced psychopathologies. *Biochem. Mosc.* **82:** 351–365
